# Improving the calling of non-invasive prenatal testing on 13-/18-/21-trisomy by support vector machine discrimination

**DOI:** 10.1101/216689

**Authors:** Jianfeng Yang, Xiaofan Ding, Weidong Zhu

**Keywords:** Next-generation sequencing (NGS), non-invasive prenatal testing (NIPT), Z test, support vector machine (SVM)

## Abstract

With the advance of next-generation sequencing technologies, non-invasive prenatal testing (NIPT) has been developed and employed in fetal aneuploidy screening on 13-/18-/21-trisomies through detecting cell-free fetal DNA (cffDNA) in maternal blood. Although Z test is widely used in NIPT nowadays, there is still necessity to improve its accuracy for removing a) false negatives and false positives, and b) the ratio of unclassified data, so as to reduce the potential harm to patients caused by these inaccuracies as well as the induced cost of retests.

Employing multiple Z tests with machine-learning algorithm could provide a better prediction on NIPT data. Combining the multiple Z values with indexes of clinical signs and quality control, features were collected from the known samples and scaled for model training in support vector machine (SVM) discrimination. The trained model was applied to predict the unknown samples, which showed significant improvement. In 4752 qualified NIPT data, our method reached 100% accuracies on all three chromosomes, including 151 data that were grouped as unclassified by one-Z-value based method. Moreover, four false positives and four false negatives were corrected by using this machine-learning model.

To our knowledge, this is the first study to employ support vector machine in NIPT data analysis. It is expected to replace the current one-Z-value based NIPT analysis in clinical use.

## Maintext: Introduction

Non-invasive prenatal testing (NIPT), which was based on the discovery of cell-free fetal DNA (cffDNA) in maternal plasma and serum, has been applied in clinical use for fetal aneuploidy detection mainly on Down’s syndrome, Edward’s syndrome and Patau’s syndrome, respectively corresponding to 21-trisomy, 18-trisomy and 13-trisomy [1-3]. Though the golden standard on prenatal diagnosis of fetal aneuploidy is the invasive amniocentesis that have a rate of 1/250 could lead to abortion, NIPT was sufficiently robust that the International Society for Prenatal Diagnosis [4], the National Society of Genetic Counselors [5], the American College of Obstetricians and Gynecologists and the Society for Maternal–Fetal Medicine [6] had published committee opinions, stating that cffDNA testing could be offered to pregnant women at high risk for fetal aneuploidy as a screening option after counseling.

Except those employing deep sequencing or array-based methods, most NIPT were performed using the low-coverage next-generation sequencing (NGS) platforms such as Verifi [7], Materni21 [8], panorama [9], NIFTY [10] and so on. Generally, the reads generated on the sequencing platforms mentioned above were aligned to genomics position and counted as depth in bins of a certain size. Then the depth was normalized and compared with reference control of negative samples [1] or the internal reference chromosomes to measure the deviation [11]. Due to the deviation between the normal fetal diploid and abnormal fetal triploid was as small as around 2%-5%, Z test was frequently employed in analysis to test whether the hypothesis that the value of unknown data is different from the mean of control data is accepted or not.

However, some problems still need to be solved as follows: a) only one Z value may not be sufficient to give a correct prediction to each sample due to read distribution bias in individual case; b) fetal fraction, a factor proven to be crucial in trisomy determination in NIPT, was however not involved in sample discrimination in most of methods nowadays; c) the sample with Z score inside an ambiguous interval called “grey zone”, ranging from 1.96 to 4, would be failed in prediction and hence required a retest, resulting in multiplying the cost. As shown in Figure 1 and Supplementary Figure 1, the current one-Z-value based NIPT was not able to discriminate samples inside grey zone, especially when fetal fraction is around or less than 5%. Such problems mentioned above could result in higher cost of testing and delay of appropriate treatment.

**Figure 1.**
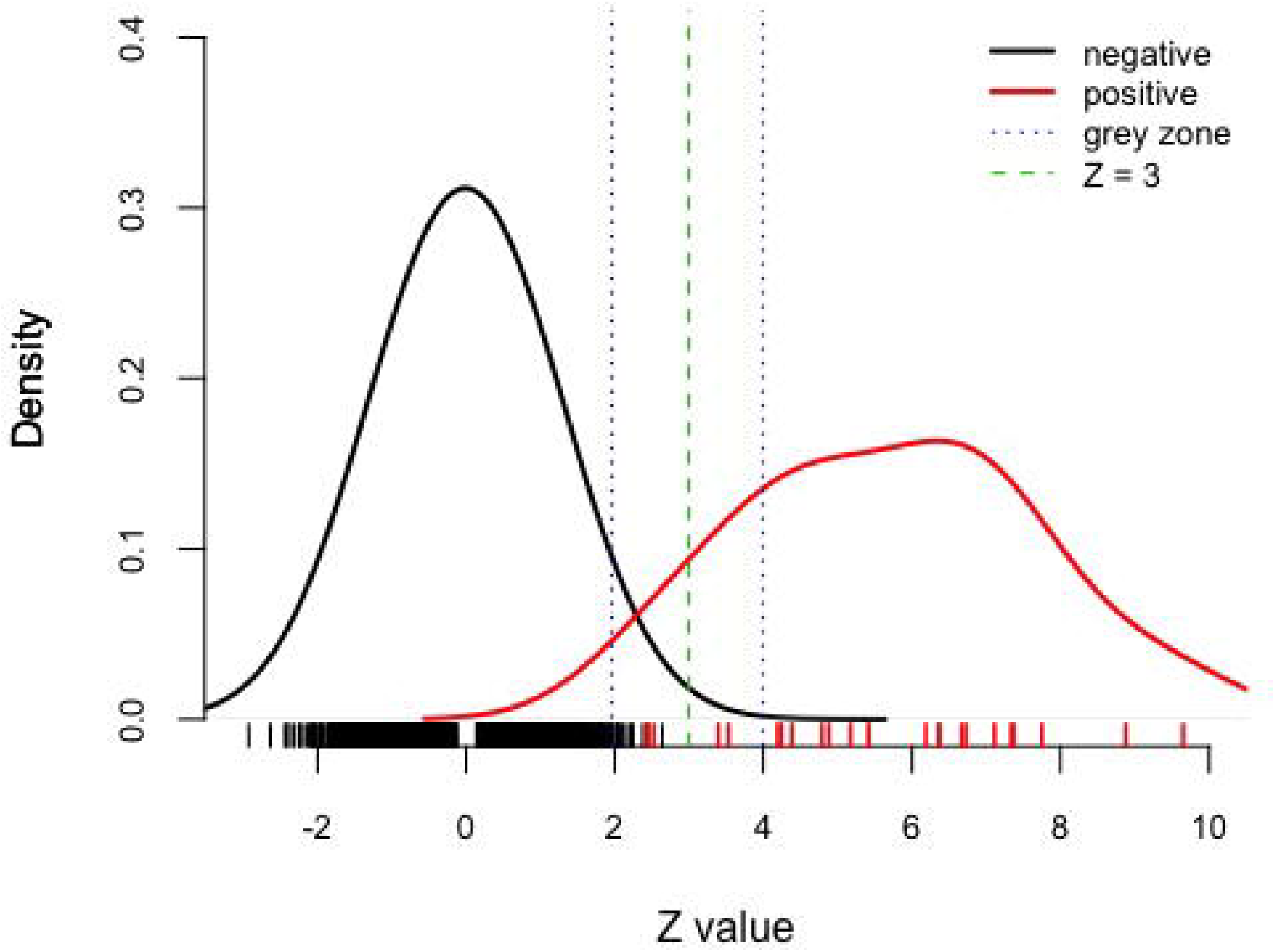
Density plot of Z values from current one-Z-test based NIPT. Negatives (only those with Z > 0 were shown) and positives are shown in dark and red respectively. Green dash indicates the cutoff of Z = 3 that was frequently used as a criterion in discrimination. Blue dashes shows the "grey zone" interval between Z = 1.96 and 4, which means failure in discrimination and requires a retest.

Therefore, it is meaningful to develop a more precise method for NIPT calling. The support vector machine (SVM) is an excellent tool for this purpose. It is a supervised machine learning algorithm that identifies an arbitrarily defined framework for discriminating query data using a model trained from selected features [12]. SVM has already shown high robustness and accuracy in fields [13], such as cancer subtype classification [14], splice site prediction [15] and single nucleotide polymorphism (SNP) prediction [16]. Considering the advantages of SVM that: 1) more features could give more accurate prediction and 2) feature co-linearity would not affect the discrimination, SVM algorithm was selected to train models for NIPT prediction in this study.

Three types of Z tests were employed in analysis: Z_baseline, Z_chromosome and Z_sample. Each type of Z test was performed twice: one with the negative control, and the other with the assumed positive control. The latter was lack in most of NIPT currently, however it is very important for removing false positives. Actually NIFTY of BGI performed student’s T tests on both negative control and assumed positive control following by a logarithmic likelihood odds ratio calculation [10], however it ignored the variances inside samples and between chromosomes as well as other clinical features. Combining multiple Z values with indexes of clinical signs and quality control, a support-vector machine algorithm was employed to train models for accurately discriminate the NIPT samples, especially the “grey zone” NIPT results as well as those falsely predicted before.

## Materials and Methods

### Specimen source

This study was based on a retrospective analysis of data prospectively generated on consecutive clinical samples for the NIPT from March to July 2016 at GuangZhou DaAn Clinical Laboratory Centre that was one of nine laboratories approved to report official NIPT result for clinical use in mainland China since late 2014.

All specimens were processed by NIPT pipeline provided by DaAn Gene Co. Ltd., using reagent kit and semi-conductor sequencing platform certified by Chinese Food and Drug Administration (CFDA). The reported results were output through a CFDA-certified standard operation protocol (SOP) and a DaAn Gene’s compiled bioinformatics plugin named “Seqboost” developed on the basis of Liao et al ‘s paper in 2013 [17] that described a one-Z-value based NIPT approach on semi-conductor sequencing platform. Since all the CFDA-certified NIPT reports were restricted on the three chromosomes 13, 18 and 21, our study would specially focus on the detection of these specified chromosomes.

### Data summary

In total 5518 NIPT data were collected during the period from two semi-conductor sequencers located in the lab in Guangzhou (See Table 1). Forty-seven of them were positives, including five for trisomy 13, fifteen for trisomy 18 and twenty-seven for trisomy 21. Average age of pregnant mother with negative results was 31.57 (95% CI: 15-51), smaller than the average age of ones with positive results (32.83, 95% CI: 17-47). Another 500 negative samples were recruited as reference negative control for
NIPT calling.

**Table 1.** Demographic Subjects of pregnant women undergoing non-invasive prenatal testing (NIPT) for aneuploidies between 1 March and 31 July in 2016

| Subject <sup>a</sup> | Total | % of all | Negative <sup>b</sup> | % in group | % of all | Positive <sup>b</sup> | % in group | % of all | P13 <sup>b</sup> | % in group | % of all | P18 <sup>b</sup> | % in group | % of all | P21 <sup>b</sup> | % in group | % of all |
| --- | --- | --- | --- | --- | --- | --- | --- | --- | --- | --- | --- | --- | --- | --- | --- | --- | --- |
|  | 5518 |  | 5473 |  | 99.18 | 45 |  | 0.82 | 5 |  | 0.09 | 14 |  | 0.25 | 27 |  | 0.49 |
| Age <sup>ac</sup> | 31.58 | (15-47) | 31.83 | (15-47) |  | 31.56 | (20-43) |  | 31.2 | (25-40) |  | 29.57 | (23-42) |  | 32.59 | (20-43) |  |
| <24 | 700 | 12.69 | 691 | 12.63 | 98.71 | 9 | 20.00 | 1.29 | 0 | 0.00 | 0.00 | 4 | 28.57 | 0.57 | 5 | 18.52 | 0.71 |
| 25-29 | 1285 | 23.29 | 1273 | 23.26 | 99.07 | 12 | 26.67 | 0.93 | 3 | 60.00 | 0.23 | 5 | 35.71 | 0.39 | 4 | 14.81 | 0.31 |
| 30-34 | 1371 | 24.85 | 1368 | 25.00 | 99.78 | 3 | 6.67 | 0.22 | 0 | 0.00 | 0.00 | 1 | 7.14 | 0.07 | 3 | 11.11 | 0.22 |
| 35-40 | 1741 | 31.55 | 1727 | 31.55 | 99.20 | 14 | 31.11 | 0.80 | 1 | 20.00 | 0.06 | 1 | 7.14 | 0.06 | 12 | 44.44 | 0.69 |
| >40 | 421 | 7.63 | 414 | 7.56 | 98.34 | 7 | 15.56 | 1.66 | 1 | 20.00 | 0.24 | 3 | 21.43 | 0.71 | 3 | 11.11 | 0.71 |
| Week <sup>ac</sup> | 17.4 | (8-37) | 17.2 | (8-37) |  | 15.91 | (12-21) |  | 13.6 | (12-15) |  | 16.21 | (12-20) |  | 16.33 | (12-21) |  |
| <13 | 651 | 11.80 | 643 | 11.75 | 98.77 | 8 | 17.78 | 1.23 | 2 | 40.00 | 0.31 | 1 | 7.14 | 0.15 | 5 | 18.52 | 0.77 |
| 14-27 | 4807 | 87.11 | 4770 | 87.16 | 99.23 | 37 | 82.22 | 0.77 | 3 | 60.00 | 0.06 | 13 | 92.86 | 0.27 | 22 | 81.48 | 0.46 |
| >28 | 60 | 1.09 | 60 | 1.10 | 100.00 | 0 | 0.00 | 0.00 | 0 | 0.00 | 0.00 | 0 | 0.00 | 0.00 | 0 | 0.00 | 0.00 |
| CostDay <sup>ac</sup> | 10.39 | (5-62) | 10.38 | (5-62) |  | 10.67 | (6-18) |  | 10 | (8-14) |  | 12.07 | (7-18) |  | 10.11 | (6-17) |  |
| <7 | 956 | 17.33 | 947 | 17.30 | 99.06 | 9 | 20.00 | 0.94 | 0 | 0.00 | 0.00 | 2 | 14.29 | 0.21 | 7 | 25.93 | 0.73 |
| 8-14 | 3963 | 71.82 | 3933 | 71.86 | 99.24 | 30 | 66.67 | 0.76 | 5 | 100.00 | 0.13 | 9 | 64.29 | 0.23 | 17 | 62.96 | 0.43 |
| 15-21 | 561 | 10.17 | 555 | 10.14 | 98.93 | 6 | 13.33 | 1.07 | 0 | 0.00 | 0.00 | 3 | 21.43 | 0.53 | 3 | 11.11 | 0.53 |
| >22 | 38 | 0.69 | 38 | 0.69 | 100.00 | 0 | 0.00 | 0.00 | 0 | 0.00 | 0.00 | 0 | 0.00 | 0.00 | 0 | 0.00 | 0.00 |

<sup>a</sup> Age means the age of the pregnant mother while doing the NIPT; Week means the gestational week while doing the NIPT; CostDay means the time cost in our NIPT service.
<sup>b</sup> Positive means the trisomy in either chromosome 13, 18 or 21. If none of these three chromosomes were found trisomy, the sample would be regarded as Negative in this study. P13 means trisomy in chromosome 13; P18 means trisomy in chromosome 18; P21 means trisomy in chromosome 21.
<sup>c</sup> Average values of relevant subjects with minimums and maximums in the brackets

As shown in Supplementary Table 1, a series of values were listed to demonstrate the information of these data, including “Z_run” as the Z scores output by Seqboost in one’s run, “Real_state” as the results confirmed by prenatal or postnatal diagnosis, fetal fraction predicted using SeqFF [18], peak value of read length, maternal age and gestational week. According to CFDA’s NIPT policy and DaAn Gene’s SOP, Z score = 3 is the cutoff to distinguish negatives and positives. Hence in routine NIPT, the data with “Z_run > 3” would be regarded as positive, meaning it’s significantly deviated from the baseline of reference dataset; while those with “Z_run ≥< 3” would be regarded as negative. Hence, the data predicted as positive with “Real_state = −1” as negative were false positives; those predicted as negative with “Real_state = 1” as positive were false negatives.

Of these 5518 data, 766 data with unique reads fewer than 3,000,000 or predicted fetal fraction less than 5% were labeled as “QC-filtered” on the basis of quality control (QC) according to the SOP. The remaining QC-pass 4752 data were categorized into three groups for specified chromosomes on the basis of the principle of statistics: Group “N” as those with Z scores smaller than 1.96, meaning not significant higher than baseline of reference dataset (p > 0.05); Group “P” as those with Z scores larger than 4, meaning significant higher than baseline of reference dataset (p < 0.0001); Group “Unclassified” as those with Z scores between 1.96 and 4, meaning retest is required for double check in nowadays’ NIPT. For each specified chromosome, data in Groups “N” and “P” were employed to train models and conduct internal validation in this study. Data in Group “Unclassified” and “QC-filtered” were used in performance test, as well as the six false positives and nine false negatives happened in previous NIPT reports.

### Feature selection and data reanalysis

Reads generated from semi-conductor sequencer were already trimmed and mapped to hg19, following by recalibration and realignment through Ion Torrent Suite Software. Then unique reads were obtained by using ‘samtools view –F 1024 –q 10’ [19]. Similar with the CFDA-certified DaAn Gene’s SOP, read-depth for each contiguous 20kb bin was calculated using the genomeCoverageBed program in BEDtools [20]. To remove the bias of read-depth distribution caused by data volume difference, GC content and casual sequencing bias respectively, three types of normalization were applied in four steps: 1) Intra-run normalization was used to eliminate the difference between each data; 2) Winsorization that was a transformation reducing the influence of outliers by moving observations outside a certain fractile in the distribution to that fractile [21], was employed to reduce the extreme read-depth among each contiguous window consisting of 15 bins of 20 kb; 3) LOESS was employed to remove GC-bias as written in Chiu’s paper [1]; 4) Intra-run normalization again due to steps 2) and 3) could induce bias of data size. Mean and standard deviation (s.d.) of read-depth of each chromosome were calculated for further statistic analysis.

The normalized read-depth of each bin was added up every 15 bins to smooth the read-depth signal. Then the mean and standard deviation of merged read-depth on each chromosome was calculated to statistic analysis for fetal aneuploidy evaluation.

For each data, six Z scores were called as described by the following formula:

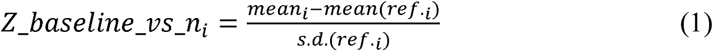

where *Z_baseline_vs_n* means the Z score normalized to the average of reference negative samples on chromosome *i*, and *ref.* means the normalized read-depth values of reference negative samples.

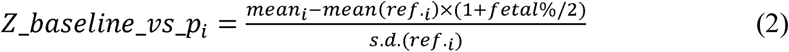

where *Z_baseline_vs_p* means the Z score normalized to the average of predicted reference positive data, *fetal%* means fetal DNA fraction. The predicted reference positive data is equal to the mean value of reference negative data multiplied by a factor ‘1+fetal%/2’ based on the assumption that half of fetal fraction would be increased when trisomy happens.

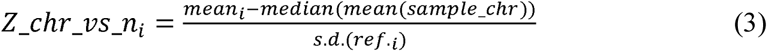

where *Z_chr_vs_n* means the Z score normalized to the internal reference autosome value that is the median of all averages of normalized read-depth in each autosome of this sample, which was similar in Lau’s paper [11].

Similarly, we have:

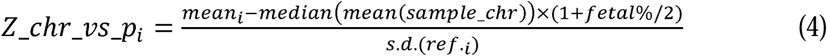

where *Z_chr_vs_p* means the Z score normalized to the predicted positive internal reference autosome value that is the median of predicted positive averages of normalized read-depth in each autosome of this sample.

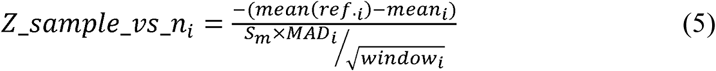

where *Z_sample_vs_n* means the Z score normalized to the average of sample data, *MAD* means the median absolute deviation of read-depth, *window* means the number of windows on the chromosome *i*, and *Sm* is a factor equal to 1.4826 and makes 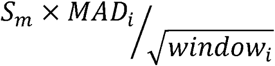 approximate to the standard deviation of read-depth of sample data.

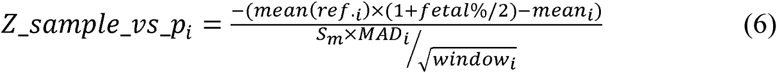

where Z_sample_vs_p_i_ means the Z score normalized to the mean value of predicted positive sample data.

### SVM discrimination

Six Z values together with fetal fraction, peak value of read length, maternal age and gestational week, were collected for support vector machine classification. For the ten features selected for SVM classification model training, the six Z score-based features were essential because their distributions between negatives and positives were significant different (p < 2.2×10^−16^), while the other four features were not significant biased (See Table 2).

**Table 2.** List of features employed in SVM classification

| Feature Number | Feature Name | Description | SVM Scale | *p*-value <sup>e</sup> |
| --- | --- | --- | --- | --- |
| D1 | Z_baseline_vs_n | Z value normalized to the baseline of control samples | No | $< 2.2e^{-16}$ |
| D2 | Z_baseline_vs_p | Z value normalized to the baseline of predictive positive samples | No | $< 2.2e^{-16}$ |
| D3 | Z_chr_vs_n | Z value normalized to the internal chromosome reference | No | $< 2.2e^{-16}$ |
| D4 | Z_chr_vs_p | Z value normalized to the predictive positive internal chromosome | No | $< 2.2e^{-16}$ |
| D5 | Z_sample_vs_n | Z value normalized to the baseline of control samples | No | $< 2.2e^{-16}$ |
| D6 | Z_sample_vs_p | Z value normalized to the baseline of predictive positive samples | No | $< 2.2e^{-16}$ |
| D7 | Fetal | Fetal fraction in maternal plasma | Yes | 0.7542 |
| D8 | Peak | Peak value of read length distribution | Yes | 0.6655 |
| D9 | MA | Maternal age | Yes | 0.2541 |
| D10 | GW | Gestational week | Yes | 0.5125 |

<sup>e</sup> Wilcoxon rank-sum test

Libsvm package [12] was employed to achieve SVM discrimination in this study. As described in the proposed SVM workflow (Figure 2), ten features were collected from the data in Groups “N” and “P” for model building on specified chromosomes. The six Z scores obtained from formula (1) to (6) do not need scaling due to they were already normalized, while the other 4 features including fetal fraction, peak value of read length, maternal age and gestational week, would be normalized to same scale ranging from 0 to 3 by the command ‘svm-scale –l 0 –u +3’. For the known data, ‘-s’ was used to save the scaling range, while ‘-r’ was used to restore the saved scaling range on unknown data. Then, the SVM model was constructed by ‘svm-train’, using SVM type ‘c-SVC’ for multiple classification and kernel type ‘RBF (radial basis function)’ for non-linear SVM model in default. This model was employed to do prediction by ‘svm-predict’ in Libsvm package.

**Figure 2.**
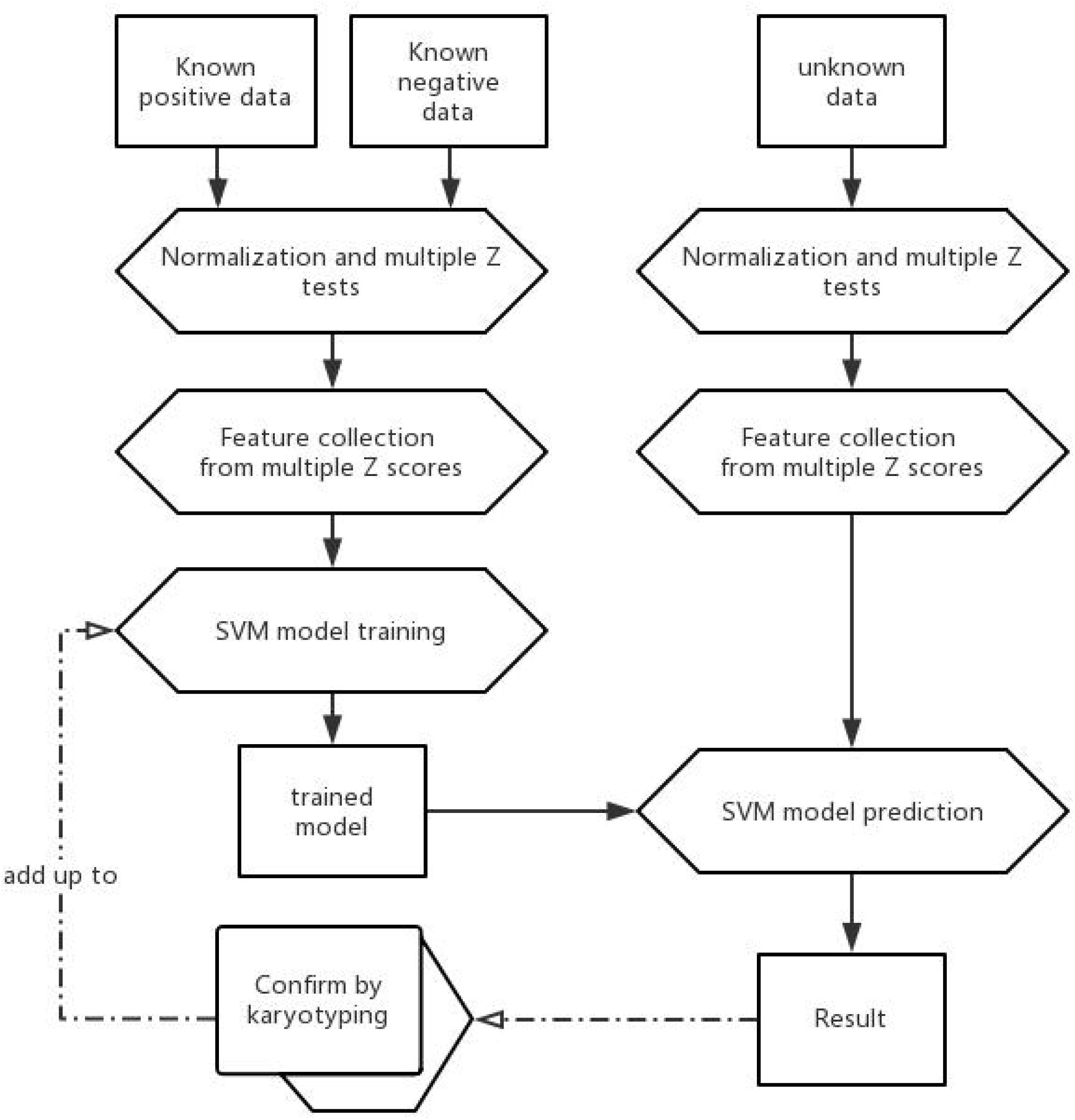
Pipeline of SVM-based NIPT in this study. The SVM model was trained from known data of both negatives and positives. The predicted results could be added up to known dataset for model rebuild after confirm by karotyping.

### Performance test of SVM classification model in predicting NIPT data

Firstly for the data in Groups “N” and “P” on specified chromosomes, an internal validation was done using the model built based on these data themselves. Importantly, the trained models were applied to predict the data in Group “Unclassified”, which was the most meaningful application in this study. As well, the models were applied to predict the data in Group “QC-filtered”.

### Comparison with other discrimination methods

Other discrimination methods such as linear discriminant analysis (LDA), quadratic discriminant analysis (QDA), decision tree (Dtree) and Neuron network (Nnet) were also tested on the same NIPT dataset in this study. An R package ‘MASS’ was applied to test LDA and QDA models built from the selected ten features. For Dtree and Nnet, R packages ‘party’ and ‘nnet’ were applied in this comparison of performance tests. Similar to the above statistic on the performance test of SVM models, the statistics of these four discrimination models were performed on three groups of data respectively:

1) Group “N” and “P”; 2) Group “Unclassified”; 3) Group “QC-filtered”. For visualization of comparison, models built from two of the ten features by the five discrimination methods were tested, using feature 1 from formula (1) and feature 3 from formula (3). The two-dimension hyper-planes for discrimination were plotted using ‘contour’ in R package ‘graphic’.

## Results

### Visualization of discrimination tools on two dimensions

As an example shown in Figure 3, nearly all of five types of discrimination models illustrated good classification lines on the trained dataset (Groups “N” and “P”) using two out of ten features, except LDA that was with one false negative. The two features D1 and D3 were obtained from formula (1) and (3), representing two Z values normalized to different references (See Materials and Methods). A 3-D plot and its three 2-D plots were also given in Supplementary Figure 2 to show how SVM model works in discriminating negatives and positives. These results were for visualization of how discrimination tools separate the data, whereas in reality all of ten features would be employed.

**Figure 3.**
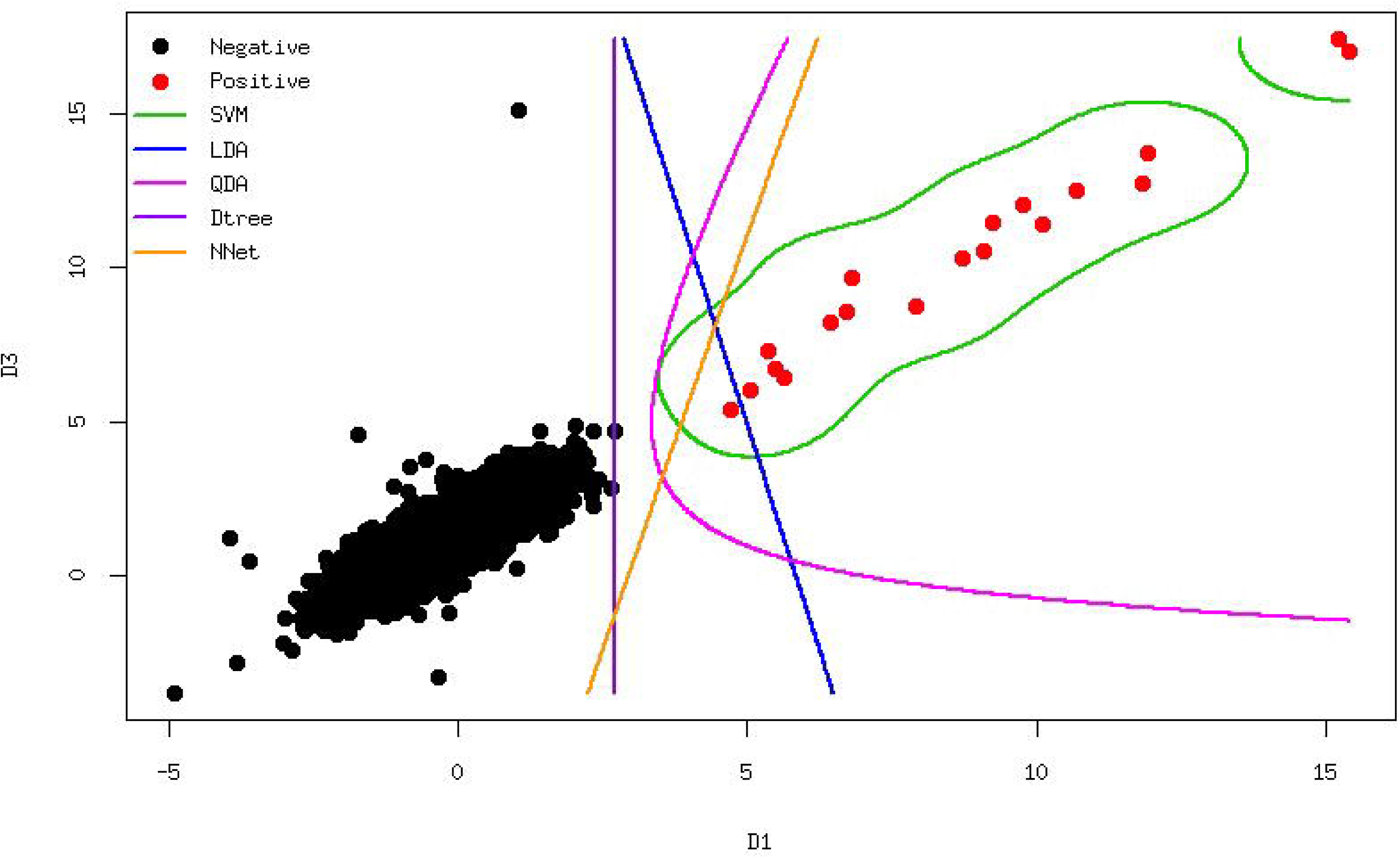
A 2-D contour plot of five discrimination models on NIPT data of Group "N" and "P" on chromosome 21. Features D1 and D3 were applied in this visualization and represented as X-axis and Y-axis respectively. Dark solid points illustrate the negative samples and red solid points the positive samples. The five two-dimension hyper-planes for discrimination (green for SVM, blue for LDA, pink for QDA, purple for decision tree and orange for neuron network) were drawn on the basis of predicted categories, using 'contour' in R package 'graphic'.

### Performance of SVM classification models

Table 3 demonstrated the performances of different discrimination models on NIPT prediction. As internal validations, the SVM models predict the training data with 100% accuracy on all three chromosomes. For chromosome 21, 4691 data were employed in model training, including 19 positives (Z score >=4) and 4672 negatives (Z score <= 1.96). Of these 4691 data, 134 were effective as support vectors in model training, including 16 positives and 118 negatives. For chromosome 18, 4704 data were employed in model training, including 7 positives and 4697 negatives. Of these 4704 data, 189 were effective as support vectors in model training, including 7 positives and 182 negatives. For chromosome 13, 4710 data were employed in model training, including 4 positives and 4706 negatives. Of these 4710 data, 211 were effective as support vectors in model training, including 4 positives and 207 negatives.

**Table 3.**
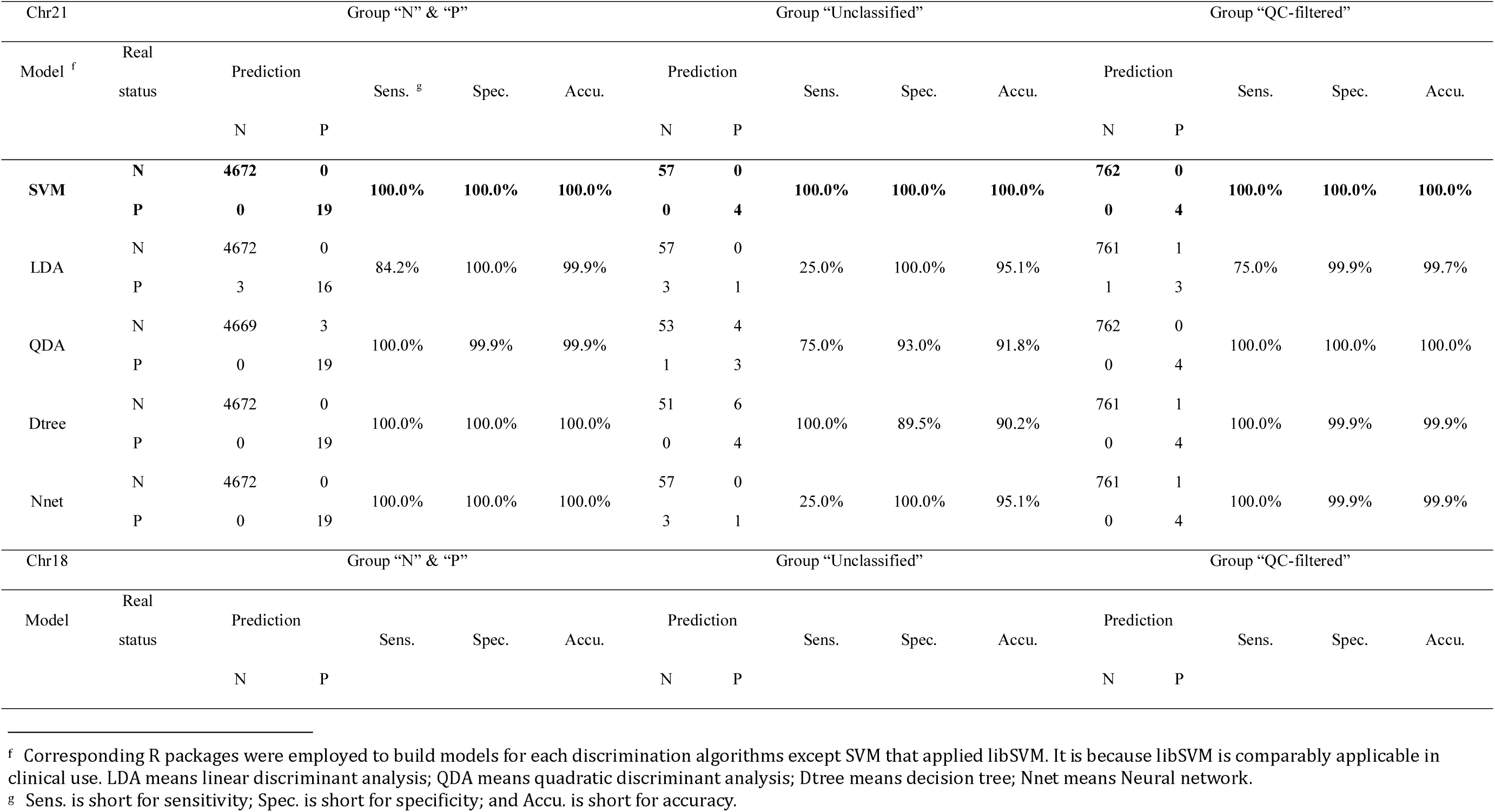
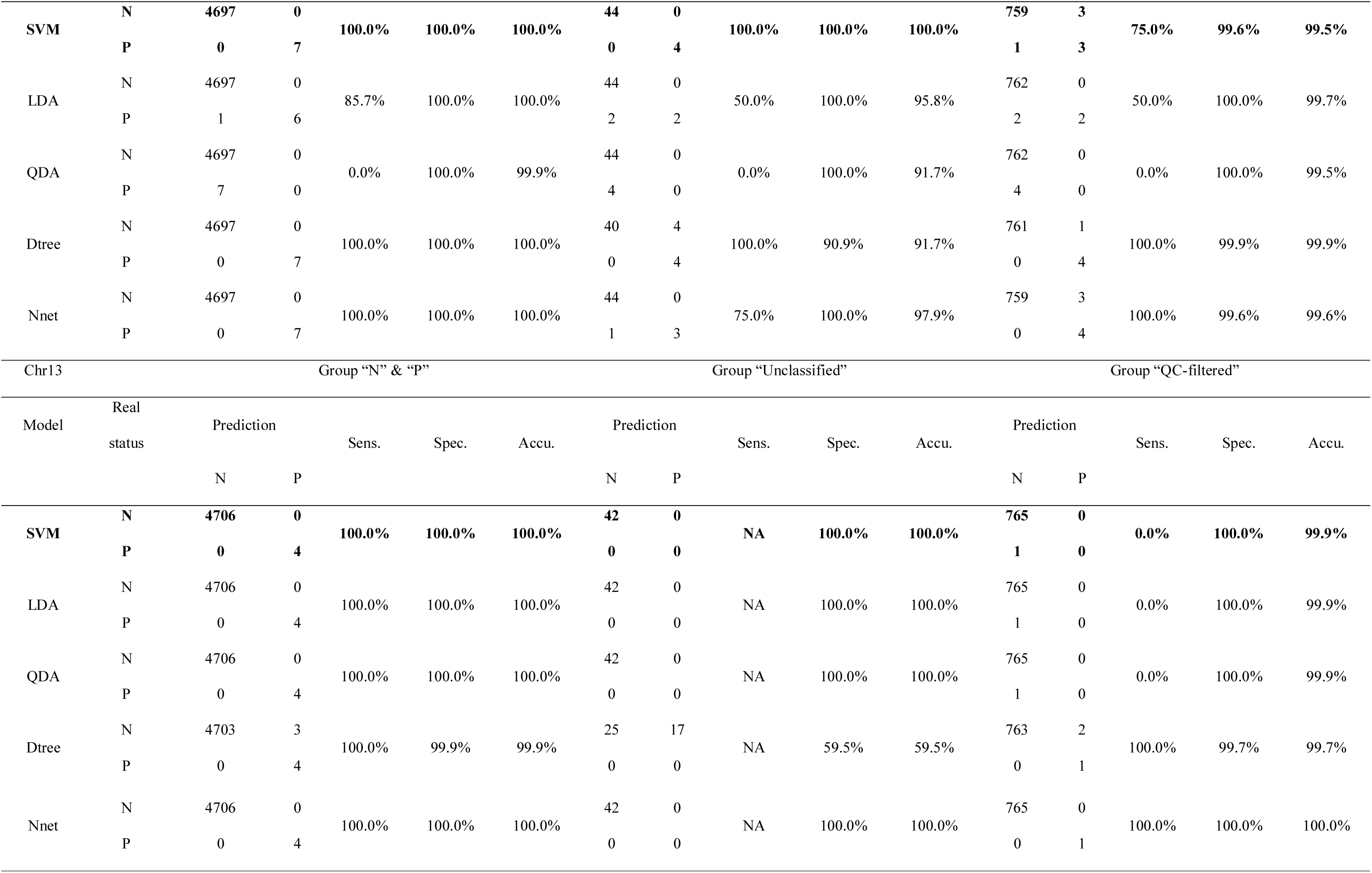
Performance of different discrimination models on NIPT prediction using ten selected features

| Chr21 |  | Group “N” & “P” |  |  |  |  | Group “Unclassified” |  |  |  |  | Group “QC-filtered” |  |  |  |  |
| --- | --- | --- | --- | --- | --- | --- | --- | --- | --- | --- | --- | --- | --- | --- | --- | --- |
| Model <sup>f</sup> | Real status | Prediction |  | Sens. <sup>g</sup> | Spec. | Accu. | Prediction |  | Sens. | Spec. | Accu. | Prediction |  | Sens. | Spec. | Accu. |
|  |  | N | P |  |  |  | N | P |  |  |  | N | P |  |  |  |
| SVM | N | **4672** | **0** | **100.0%** | **100.0%** | **100.0%** | **57** | **0** | **100.0%** | **100.0%** | **100.0%** | **762** | **0** | **100.0%** | **100.0%** | **100.0%** |
|  | P | **0** | **19** |  |  |  | **0** | **4** |  |  |  | **0** | **4** |  |  |  |
| LDA | N | 4672 | 0 | 84.2% | 100.0% | 99.9% | 57 | 0 | 25.0% | 100.0% | 95.1% | 761 | 1 | 75.0% | 99.9% | 99.7% |
|  | P | 3 | 16 |  |  |  | 3 | 1 |  |  |  | 1 | 3 |  |  |  |
| QDA | N | 4669 | 3 | 100.0% | 99.9% | 99.9% | 53 | 4 | 75.0% | 93.0% | 91.8% | 762 | 0 | 100.0% | 100.0% | 100.0% |
|  | P | 0 | 19 |  |  |  | 1 | 3 |  |  |  | 0 | 4 |  |  |  |
| Dtree | N | 4672 | 0 | 100.0% | 100.0% | 100.0% | 51 | 6 | 100.0% | 89.5% | 90.2% | 761 | 1 | 100.0% | 99.9% | 99.9% |
|  | P | 0 | 19 |  |  |  | 0 | 4 |  |  |  | 0 | 4 |  |  |  |
| Nnet | N | 4672 | 0 | 100.0% | 100.0% | 100.0% | 57 | 0 | 25.0% | 100.0% | 95.1% | 761 | 1 | 100.0% | 99.9% | 99.9% |
|  | P | 0 | 19 |  |  |  | 3 | 1 |  |  |  | 0 | 4 |  |  |  |
| Chr18 |  | Group “N” & “P” |  |  |  |  | Group “Unclassified” |  |  |  |  | Group “QC-filtered” |  |  |  |  |
| Model | Real status | Prediction |  | Sens. | Spec. | Accu. | Prediction |  | Sens. | Spec. | Accu. | Prediction |  | Sens. | Spec. | Accu. |
|  |  | N | P |  |  |  | N | P |  |  |  | N | P |  |  |  |

<sup>f</sup> Corresponding R packages were employed to build models for each discrimination algorithms except SVM that applied libSVM. It is because libSVM is comparably applicable in clinical use. LDA means linear discriminant analysis; QDA means quadratic discriminant analysis; Dtree means decision tree; Nnet means Neural network.
<sup>g</sup> Sens. is short for sensitivity; Spec. is short for specificity; and Accu. is short for accuracy.

|  |  |  |  |  |  |  |  |  |  |  |  |  |  |  |  |  |
| --- | --- | --- | --- | --- | --- | --- | --- | --- | --- | --- | --- | --- | --- | --- | --- | --- |
| SVM | N | 4697 | 0 | 100.0% | 100.0% | 100.0% | 44 | 0 | 100.0% | 100.0% | 100.0% | 759 | 3 | 75.0% | 99.6% | 99.5% |
|  | P | 0 | 7 |  |  |  | 0 | 4 |  |  |  | 1 | 3 |  |  |  |
| LDA | N | 4697 | 0 | 85.7% | 100.0% | 100.0% | 44 | 0 | 50.0% | 100.0% | 95.8% | 762 | 0 | 50.0% | 100.0% | 99.7% |
|  | P | 1 | 6 |  |  |  | 2 | 2 |  |  |  | 2 | 2 |  |  |  |
| QDA | N | 4697 | 0 | 0.0% | 100.0% | 99.9% | 44 | 0 | 0.0% | 100.0% | 91.7% | 762 | 0 | 0.0% | 100.0% | 99.5% |
|  | P | 7 | 0 |  |  |  | 4 | 0 |  |  |  | 4 | 0 |  |  |  |
| Dtree | N | 4697 | 0 | 100.0% | 100.0% | 100.0% | 40 | 4 | 100.0% | 90.9% | 91.7% | 761 | 1 | 100.0% | 99.9% | 99.9% |
|  | P | 0 | 7 |  |  |  | 0 | 4 |  |  |  | 0 | 4 |  |  |  |
| Nnet | N | 4697 | 0 | 100.0% | 100.0% | 100.0% | 44 | 0 | 75.0% | 100.0% | 97.9% | 759 | 3 | 100.0% | 99.6% | 99.6% |
|  | P | 0 | 7 |  |  |  | 1 | 3 |  |  |  | 0 | 4 |  |  |  |
| Chr13 |  | Group “N” & “P” |  |  |  |  |  | Group “Unclassified” |  |  |  |  |  | Group “QC-filtered” |  |  |
| Model | Real status | Prediction |  | Sens. | Spec. | Accu. | Prediction |  | Sens. | Spec. | Accu. | Prediction |  | Sens. | Spec. | Accu. |
|  |  | N | P |  |  |  | N | P |  |  |  | N | P |  |  |  |
| SVM | N | 4706 | 0 | 100.0% | 100.0% | 100.0% | 42 | 0 | NA | 100.0% | 100.0% | 765 | 0 | 0.0% | 100.0% | 99.9% |
|  | P | 0 | 4 |  |  |  | 0 | 0 |  |  |  | 1 | 0 |  |  |  |
| LDA | N | 4706 | 0 | 100.0% | 100.0% | 100.0% | 42 | 0 | NA | 100.0% | 100.0% | 765 | 0 | 0.0% | 100.0% | 99.9% |
|  | P | 0 | 4 |  |  |  | 0 | 0 |  |  |  | 1 | 0 |  |  |  |
| QDA | N | 4706 | 0 | 100.0% | 100.0% | 100.0% | 42 | 0 | NA | 100.0% | 100.0% | 765 | 0 | 0.0% | 100.0% | 99.9% |
|  | P | 0 | 4 |  |  |  | 0 | 0 |  |  |  | 1 | 0 |  |  |  |
| Dtree | N | 4703 | 3 | 100.0% | 99.9% | 99.9% | 25 | 17 | NA | 59.5% | 59.5% | 763 | 2 | 100.0% | 99.7% | 99.7% |
|  | P | 0 | 4 |  |  |  | 0 | 0 |  |  |  | 0 | 1 |  |  |  |
| Nnet | N | 4706 | 0 |  |  |  | 42 | 0 |  |  |  | 765 | 0 |  |  |  |
|  | P | 0 | 4 | 100.0% | 100.0% | 100.0% | 0 | 0 | NA | 100.0% | 100.0% | 0 | 1 | 100.0% | 100.0% | 100.0% |

Importantly, the SVM classification model was employed to predict the other QC-pass data used to be regarded as “unclassified” (1.96< Z score < 4), which gave us a 100% accuracy prediction result. For chromosome 21, all 61 data in “grey zone” (1.3% of all QC-pass data) were accurately predicted using the training model, including 4 positives and 57 negatives. Actually, two of those 4 positives were wrongly regarded as “negative” before due to their Z scores were smaller than 3 (2.44 and 2.52 respectively), however, the SVM classification model could predict them correctly. This suggested that some positives may be covered if only using Z score = 3 as classification criteria, and SVM classification model could uncover such critical positives by training the known data. For chromosome 18, all 48 data in “grey zone” (1.0% of all QC-pass data) were accurately predicted using the training model, including 4 positives and 44 negatives. For chromosome 13, all 42 data in “grey zone” (0.9% of all QC-pass data) were accurately predicted as “negative” using the training model. In summary, all of the data could not be judged using only Z score (nearly 3% of all QC-pass data), were precisely predicted using the SVM algorithm with training the known data. This suggested that using SVM classification model could save around 3% of resource in retests.

Surprisingly, the SVM classification model also acted effectively in predicting the 766 QC-filtered data. For chromosome 21, the model precisely predicted all the QC-filtered data, including 4 positives and 762 negatives. For chromosome 18, 762 out of 766 data were correct (99.48%). One positive was wrongly predicted as “negative” with prediction probability 52%, while three negatives were incorrectly predicted as “positive” with predicted probabilities 50%, 92% and 99% respectively. For chromosome 13, 765 out of 766 data were correct (99.87%). Only one positive data that was regarded as “negative” with Z score = 2.79, was also incorrectly predicted as “negative” by the SVM model. This demonstrated that the SVM model could perform well in most of QC-filtered data but could not uncover all false negatives, suggesting that quality control is still necessary to guarantee the accuracy of NIPT.

### Comparison with other discrimination models in performance

Compared with other models, SVM is the only one to obtain 100% accuracy in both internal validation and prediction on data in grey zone (see Table 3). Both SVM and Nnet models obtained 100% accuracies in internal validation across three specified chromosomes. However, Nnet model did not give perfect predictions on the samples in grey zone, having three false negatives on chromosome 21 and one false negative on chromosome 18. The other three models did not perform well enough in internal validation or grey zone data prediction. It was shown that Dtree model was prone to have more false positives than false negatives in predicting. This suggested that SVM could be the best algorithm to improve the accuracy of nowadays’ NIPT.

### Application in correcting previous false callings from one-Z-test based method

In addition, we tested the model with four false positives and four false positives happened in previous time. These eight samples were wrongly predicted by Z score-based method. As shown in Table 4, all of eight samples were corrected by using the SVM classification model, according to the prediction probabilities. For the four false positives reported previously, values of features D1, D3 and D5 (three types of Z scores normalized to reference, see Methods) significantly exceeded 3, while values of features D2, D4 and D6 (three types of Z scores normalized to the predicted positives) also significantly lower than −3. This suggests that one-Z-test based method is not reliable in this situation. However, SVM model using multiple features could accurately predict the results. Besides, the SVM model performed well in correcting false negatives on chromosomes 18 and 21. Similarly, the four false negatives showed ambiguous values among features D1 to D6, suggesting that none of these six Z values could be reliable to do prediction independently. This further demonstrated that the SVM algorithm was better than the commonly used Z score-based classification approach.

**Table 4.** Correction of previous false negatives and false positives by current SVM model

| Sample | Error type <sup>h</sup> | Reported<br>Z value | D1 <sup>i</sup> | D2 | D3 | D4 | D5 | D6 | D7 | D8 | D9 | D10 | Probability<br>of negative in<br>SVM<br>prediction | Probability<br>of positive in<br>SVM<br>prediction |
| --- | --- | --- | --- | --- | --- | --- | --- | --- | --- | --- | --- | --- | --- | --- |
| chr13 |  |  |  |  |  |  |  |  |  |  |  |  |  |  |
| 13_FP_1 | FP | 3.61 | 7.80908 | -18.4017 | 10.4249 | -15.4931 | 8.12209 | -19.1393 | 22.38411066 | 151.515 | 32 | 17 | 0.998877 | 0.00112284 |
| 13_FP_2 | FP | 4.78 | 7.74425 | -9.47269 | 9.7306 | -7.34031 | 8.2322 | -10.0696 | 14.70334619 | 130.36 | 25 | 17 | 0.998162 | 0.00183793 |
| chr18 |  |  |  |  |  |  |  |  |  |  |  |  |  |  |
| 18_FN_1 | FN | 1.77 | 3.31662 | -2.6893 | 5.82699 | -0.108174 | 3.20528 | -2.59902 | 5.63756896 | 119.412 | 42 | 17 | 1.71E-007 | 1 |
| 18_FN_2 | FN | 1.43 | 5.54394 | -3.20759 | 8.52815 | -0.100806 | 5.79965 | -3.35554 | 8.214781802 | 149.353 | 41 | 16 | 1.00E-007 | 1 |
| 18_FP_1 | FP | 3.22 | 3.36731 | -10.1558 | 4.56993 | -8.8769 | 3.31148 | -9.98747 | 12.69375338 | 153.482 | 26 | 17 | 0.984857 | 0.015143 |
| 18_FN_3 | FN | 1.93 | 5.92865 | -4.21056 | 6.68878 | -3.41427 | 4.61288 | -3.2761 | 15.25077553 | 148.944 | 27 | 18 | 2.13E-005 | 0.999979 |
| chr21 |  |  |  |  |  |  |  |  |  |  |  |  |  |  |
| 21_FP_1 | FP | 2.27 | 2.17527 | -10.4262 | 4.38166 | -8.02835 | 2.35229 | -11.2747 | 17.3586114 | 139.857 | 25 | 17 | 0.997866 | 0.00213356 |
| 21_FN_1 | FN | 1.76 | 6.7041 | -3.30991 | 8.41899 | -1.47674 | 5.21281 | -2.57365 | 13.79432935 | 150.056 | 36 | 12 | 0.00435514 | 0.995645 |

<sup>h</sup> FP means false positive and FN means false negative.
<sup>i</sup> The definitions of features D1 to D10 were given in Table 2.

In summary, SVM has demonstrated its excellent performance in discrimination of NIPT results in this study, especially compared with the current one-Z-value based method. Our technique has higher sensitivity and specificity than all previously reported approaches for the detection of chromosome 13/18/21 trisomies. With this improvement, it is expected to reduce the cost of retests on samples in grey zone as well as the cost caused by false positives and false negatives. As shown in Figure 2, we expect that the SVM model could be further improved if 1) more known data were validated and added up to model-training; 2) more impacted features were discovered and added up to model-training. Some other clinical signs such as the values from serological test could be employed together with NIPT data to do prediction.

## Discussion

Our study has shown that the SVM discrimination model trained by known data could precisely predict the results for those three chromosomes in NIPT, especially for the QC-pass data. Compared with the early standard Z test-based NIPT approaches like Chiu et al ‘s [1], Chen et al ‘s [3] and Liao et al ‘s [17], our method considered fetal fraction, a factor proven to be important in NIPT analysis recently [18, 22, 23], in Z tests. BGI’s NIFTY employed a logarithmic likelihood odds ratio between binary hypotheses that took fetal fraction in consideration [10], and Yu et al improved the count-based analysis by adding another size-based approach [24]. However, these two approaches were based on one or two statistic values, such as Z > 3 [1] or L > 1 [10] to determine a sample as negative or positive. In fact, other information such as maternal age had been employed to correct bias of NIPT prediction [25]. In our SVM-based method, discrimination criterion was obtained using more information comparing with other existing NIPT algorithms.

As shown in Figure 3 and Table 3, SVM models performed better than other on these NIPT datasets. For LDA and QDA, co-linearity between features could be one of reasons of lower accuracy in prediction, while SVM allows co-linearity between features. Both decision tree and neural network performed well in internal validation but not robust enough in prediction of data in grey zone. Nevertheless, it is still worth to keep testing these machine-learning algorithms if there are more features and more data in future, since our objective is to find the best approach for clinical use.

Temporarily, SVM showed the most robustness according to this study. For the ten features selected for current SVM model training, the four non-Z-value features actually were not significantly biased in distributions between negatives and positives, though IONA’s paper reported that maternal age was useful in correcting its NIPT results [25], which might be due to the differences in sample composition.

In conclusion, out study demonstrated an accurate SVM-based algorithm for trisomy detection on chromosome 21, 18 and 13, which was the first machine-learning approach using in this field to our knowledge. This machine-learning approach could be applied in detection of aneuploidy of other chromosomes or even micro-duplication and deletion. At this moment, sex chromosome aneuploidy screening was not included in current version of SVM-based method but would be developed if we had sufficient diagnosed cases.

## List of Abbreviations

NGS: Next-Generation Sequencing

NIPT: Non-Invasive Prenatal Testing

SVM: Support Vector Machine

RBF: Radical-Based Function

CFDA: Chinese Food and Drug Administration

DNA: Deoxyribonucleic Acid

LDA: Linear Discriminant Analysis

QDA: Quadratic Discriminant Analysis

QC: Quality Control

## Authors’ Contributions

JY, WZ and TW initiated the study. JY conceived the whole study, including developing the algorithms, editing source codes, completing the bioinformatics analysis and writing the paper. SH and LX exported the sequencing data from semi-conductor sequencers, while QL exported the NIPT reports from database. All authors have read and approved the manuscript.

## Acknowledgement

The study was supported by a Program of Science and Technology in GuangZhou to Dr. Jianfeng Yang from GuangZhou Science Technology and Innovation Commission (Grant No. 201604016136) and funded by GuangZhou DaAn Clinical Laboratory Centre of YunKang Group. We thank Ms. Dayi Zheng for managing the application of patent as well as software copyright.

---

Supplementary Figure 1. Z value distributions in simulation, showing the existing problem of current one-Z-test based NIPT. Each of the three normal distributions were simulated by bootstrapping 10,000 times for negative samples (green line), positive samples with fetal fraction 5% (cyan line) and positive samples with fetal fraction 10% (red line) respectively. Yellow dash line means Z score equal to 3. Dark dash lines show the interval of grey zone. When fetal DNA fraction is around 5% that is possible to happen in real, it became difficult to distinguish positives and negatives from samples in grey zone.

Supplementary Figure 2. 3-D contour plot and its relevant 2-D plots of SVM model on NIPT data of Group “N” and “P” on chromosome 21. Features D1, D3 and D7 were applied in this visualization and represented as X-axis, Y-axis and Z-axis respectively. Dark solid points illustrate the negative samples and red solid points the positive samples.

## References

[1] R. W. K. Chiu, K. C. A. Chan, Y. Gao, V. Y. M. Lau, W. Zheng, T. Y. Leung, C. H. F. Foo, B. Xie, N. B. Y. Tsui, and F. M. F. Lun, “Noninvasive prenatal diagnosis of fetal chromosomal aneuploidy by massively parallel genomic sequencing of DNA in maternal plasma,” Proceedings of the National Academy of Sciences of the United States of America, vol. 105, no. 51, pp. 20458–20463, 2008.

[2] H. C. Fan, Y. J. Blumenfeld, U. Chitkara, L. Hudgins, and S. R. Quake, “Noninvasive diagnosis of fetal aneuploidy by shotgun sequencing DNA from maternal blood,” Proceedings of the National Academy of Sciences, vol. 105, no. 42, pp. 16266–16271, 2008.

[3] E. Z. Chen, R. W. Chiu, H. Sun, R. Akolekar, K. A. Chan, T. Y. Leung, P. Jiang, Y. W. Zheng, F. M. Lun, and L. Y. Chan, “Noninvasive prenatal diagnosis of fetal trisomy 18 and trisomy 13 by maternal plasma DNA sequencing,” PloS one, vol. 6, no. 7, pp. e21791, 2011.

[4] P. Benn, A. Borrell, H. Cuckle, L. Dugoff, S. Gross, J. A. Johnson, R. Maymon, A. Odibo, P. Schielen, and K. Spencer, “Prenatal Detection of Down Syndrome using Massively Parallel Sequencing (MPS): a rapid response statement from a committee on behalf of the Board of the International Society for Prenatal Diagnosis, 24 October 2011,” Prenatal Diagnosis, vol. 32, no. 1, pp. 1, 2012.

[5] P. L. Devers, A. Cronister, K. E. Ormond, F. Facio, C. K. Brasington, and P. Flodman, “Noninvasive Prenatal Testing/Noninvasive Prenatal Diagnosis: the Position of the National Society of Genetic Counselors,” Journal of Genetic Counseling, vol. 22, no. 3, pp. 291, 2013.

[6] “Committee Opinion No. 545: Noninvasive Prenatal Testing for Fetal Aneuploidy,” Obstetrics & Gynecology, vol. 120, no. 6, pp. 1532–1534, 2012.

[7] P. A. Taneja, H. L. Snyder, D. F. Eileen, K. M. Kruglyak, H. M. Meredith, K. J. Curnow, and B. Sucheta, “Noninvasive prenatal testing in the general obstetric population: clinical performance and counseling considerations in over 85□000 cases†,” Prenatal Diagnosis, vol. 36, no. 3, pp. 237–243, 2015.

[8] R. M. Mccullough, E. A. Almasri, X. Guan, J. A. Geis, S. C. Hicks, A. R. Mazloom, C. Deciu, P. Oeth, A. T. Bombard, and B. Paxton, “Non-invasive prenatal chromosomal aneuploidy testing--clinical experience: 100,000 clinical samples,” vol. 9, no. 10, pp. e109173, 2014.

[9] J. C. Wang, T. Sahoo, S. Schonberg, K. A. Kopita, L. Ross, K. Patek, and C. M. Strom, “Discordant Noninvasive Prenatal Testing and Cytogenetic Results,” Obstetrical & Gynecological Survey, vol. 70, no. 7, pp. 434–436, 2015.

[10] F. Jiang, J. Ren, C. Fang, Y. Zhou, J. Xie, D. Shan, S. Yue, J. Xie, B. Yin, and S. Wen, “Noninvasive Fetal Trisomy (NIFTY) test: an advanced noninvasive prenatal diagnosis methodology for fetal autosomal and sex chromosomal aneuploidies,” Bmc Medical Genomics, vol. 5, no. 1, pp. 57, 2012.

[11] T. K. Lau, F. Chen, X. Pan, R. K. Pooh, F. Jiang, Y. Li, H. Jiang, X. Li, S. Chen, and X. Zhang, “Noninvasive prenatal diagnosis of common fetal chromosomal aneuploidies by maternal plasma DNA sequencing,” The journal of maternal-fetal & neonatal medicine: the official journal of the European Association of Perinatal Medicine, the Federation of Asia and Oceania Perinatal Societies, the International Society of Perinatal Obstetricians, vol. 25, no. 8, pp. 1370–1374, 2011.

[12] C. Chih-Chung, and L. Chih-Jen, “LIBSVM: A library for support vector machines,” ACM Trans. Intell. Syst. Technol., vol. 2, pp. 1–27, 2011.

[13] S. S. Keerthi, and C.-J. Lin, “Asymptotic behaviors of support vector machines with Gaussian kernel,” Neural computation, vol. 15, no. 7, pp. 1667–1689, 2003.

[14] Y. Lee, and C.-K. Lee, “Classification of multiple cancer types by multicategory support vector machines using gene expression data,” Bioinformatics, vol. 19, no. 9, pp. 1132–1139, 2003.

[15] A. K. Baten, B. C. Chang, S. K. Halgamuge, and J. Li, “Splice site identification using probabilistic parameters and SVM classification,” BMC bioinformatics, vol. 7, no. Suppl 5, pp. S15, 2006.

[16] B. D. O’Fallon, W. Wooderchak-Donahue, and D. K. Crockett, “A support vector machine for identification of single-nucleotide polymorphisms from next-generation sequencing data,” Bioinformatics, vol. 29, no. 11, pp. 1361–1366, 2013.

[17] C. Liao, A.-h. Yin, C.-f. Peng, F. Fu, J.-x. Yang, R. Li, Y.-y. Chen, D.-h. Luo, Y.-l. Zhang, and Y.-m. Ou, “Noninvasive prenatal diagnosis of common aneuploidies by semiconductor sequencing,” Proceedings of the National Academy of Sciences, vol. 111, no. 20, pp. 7415–7420, 2014.

[18] S. K. Kim, G. Hannum, J. Geis, J. Tynan, G. Hogg, C. Zhao, T. J. Jensen, A. R. Mazloom, P. Oeth, and M. Ehrich, “Determination of fetal DNA fraction from the plasma of pregnant women using sequence read counts,” Prenatal Diagnosis, vol. 35, no. 8, pp. 810–815, 2015.

[19] H. Li, B. Handsaker, A. Wysoker, T. Fennell, J. Ruan, N. Homer, G. Marth, G. Abecasis, and R. Durbin, “The sequence alignment/map format and SAMtools,” Bioinformatics, vol. 25, no. 16, pp. 2078–2079, 2009.

[20] A. R. Quinlan, and I. M. Hall, “BEDTools: a flexible suite of utilities for comparing genomic features,” Bioinformatics, vol. 26, no. 6, pp. 841–2, Mar 15, 2010.

[21] G. Nilsen, K. Liestøl, P. Van Loo, H. K. M. Vollan, M. B. Eide, O. M. Rueda, S.-F. Chin, R. Russell, L. O. Baumbusch, and C. Caldas, “Copynumber: Efficient algorithms for single-and multi-track copy number segmentation,” BMC genomics, vol. 13, no. 1, pp. 591, 2012.

[22] P. Jiang, K. C. Chan, G. J. Liao, Y. W. Zheng, T. Y. Leung, R. W. Chiu, Y. M. Lo, and H. Sun, “FetalQuant: deducing fractional fetal DNA concentration from massively parallel sequencing of DNA in maternal plasma,” Bioinformatics, vol. 28, no. 22, pp. 2883–90, 2012.

[23] P. Jiang, X. Peng, X. Su, K. Sun, S. C. Y. Yu, W. I. Chu, T. Y. Leung, H. Sun, R. W. K. Chiu, and Y. M. D. Lo, “FetalQuantSD: accurate quantification of fetal DNA fraction by shallow-depth sequencing of maternal plasma DNA,” vol. 1, pp. 16013, 2016.

[24] S. C. Yu, P. Jiang, K. C. Allen Chan, B. H. Faas, K. W. Choy, W. C. Leung, T. Y. Leung, Y. M. Dennis Lo, and R. W. Chiu, “Combined Count- and Size-Based Analysis of Maternal Plasma DNA for Noninvasive Prenatal Detection of Fetal Subchromosomal Aberrations Facilitates Elucidation of the Fetal and/or Maternal Origin of the Aberrations,” Clinical Chemistry, 2016.

[25] P. L. C., D. D. D., F. C., F. I., and N. K.H., “IONA test for first-trimester detection of trisomies 21, 18 and 13,” Ultrasound in Obstetrics & Gynecology, vol. 47, no. 2, pp. 184–187, 2016.

